# Spatio-temporal spawning patterns and early growth of Japanese sardine in the western North Pacific during the recent stock increase

**DOI:** 10.1101/2020.12.10.420562

**Authors:** Yohei Niino, Sho Furuichi, Yasuhiro Kamimura, Ryuji Yukami

## Abstract

Spatio-temporal patterns in spawning influence growth and survival by affecting the environment experienced by offspring during their early life stages. Therefore, identifying changes in spawning patterns can help researchers understand population dynamics and recruitment fluctuations. The Japanese sardine *Sardinops melanostictus* is a small pelagic fish that undergoes large stock fluctuations. Although spawning patterns are known to change spatially and temporally with stock abundance, little information is available on the processes underlying stock increases. This study aimed to describe changes in spawning pattern and early growth of Japanese sardine during the recent period of stock increase, and to clarify the effects of different spawning periods on offspring growth. We examined trends in egg abundance in the western North Pacific in 2004–2018 and analyzed hatch dates and growth trajectories by otolith microstructure analysis of juveniles from the Kuroshio–Oyashio transitional region (the species’ main nursery area). During the study period, the main spawning area shifted from the western to the eastern part of this region off Japan, and spawning in the eastern part roughly coincided with juvenile hatch dates. Hatch dates also shifted from mid-March at the earliest to February and early March from 2013 onwards. Although early-hatched cohorts (which originate offshore of eastern Japan) experienced slower initial growth, they likely played an important role in the recent stock increase. Successful recruitment of these cohorts may have been facilitated by factors such as early hatching and transport to the nursery, which reduces the frequency of predator encounters.

## INTRODUCTION

In fishes, spatio-temporal variations in reproductive traits, such as spawning season and spawning grounds, influence subsequent growth and survival by determining the environment that offspring experience in their early life history (Secor, 2007). Therefore, identifying changes in fish reproductive traits and their impacts on the early growth and survival of offspring will help us to better understand population dynamics and recruitment fluctuations.

Sardine stocks have experienced large-scale fluctuations all over the world. The Japanese sardine *Sardinops melanostictus* is one of the most abundant species off the coast of Japan (Lluch-Belda et al., 1989; Schwartzlose et al., 1999; Yasuda et al., 1999; Yatsu, 2019). The population distributed off the Japanese Pacific coast (the Pacific stock) accounts for a considerable percentage of Japanese sardine. The Pacific stock reached a peak abundance of over 10 million t in the 1980s, but successive recruitment failures since 1988 (Watanabe et al., 1995) reduced the population to around 100,000 t in the 2000s (Furuichi et al., 2020c; Fig. 1). Since the 2010s, the stock has been in recovery thanks to favorable recruitment and reduced fishing pressure.

**Fig. 1.**
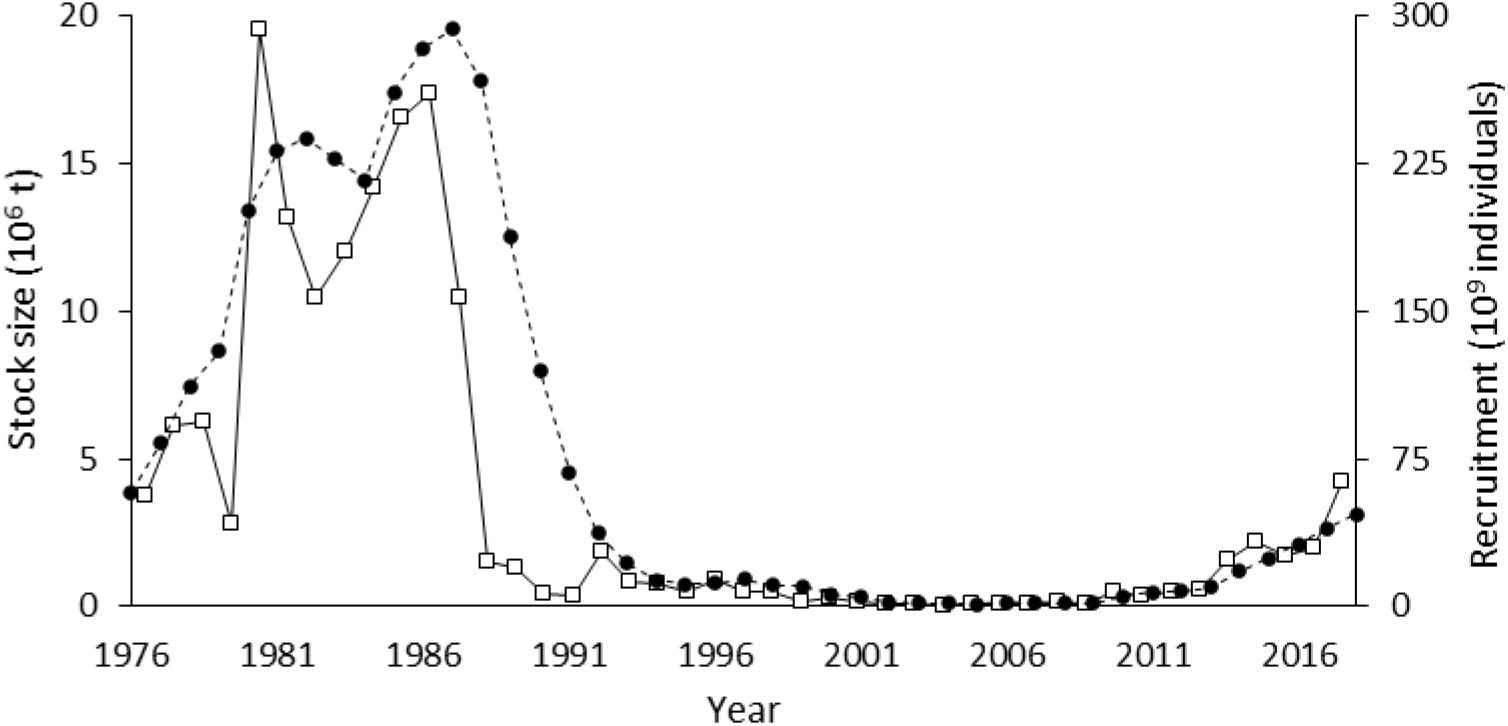
Time series of recruitment (open squares) and stock abundance (solid circles) of the Pacific stock of Japanese sardine.

The Pacific stock spawns in the Kuroshio region in winter and spring, and many eggs and larvae are transported by the Kuroshio Extension (Itoh et al., 2009; Sugisaki, 1996; Fig. 2). Juveniles begin to migrate northward as they grow and pass through their nursery area (the Kuroshio–Oyashio transitional region) during the summer and autumn (Okunishi et al., 2012; Sakamoto et al., 2019).

**Fig. 2.**
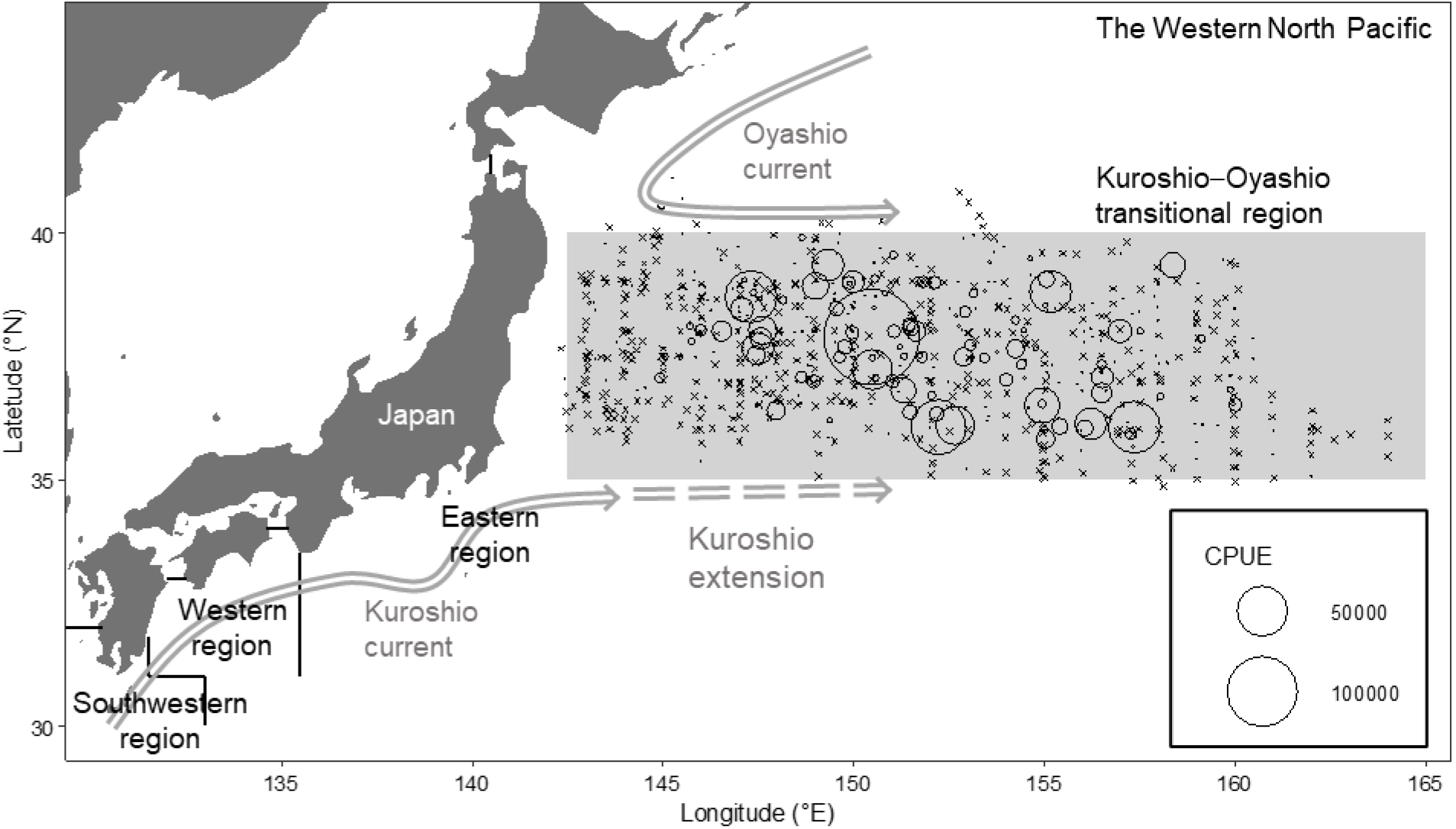
Map of the trawl survey area showing catch per unit effort (CPUE; individuals h^−1^) of juvenile Japanese sardine at each sampling station. Circle sizes indicate the magnitude of CPUE, and crosses show stations where sardine were not captured. Black lines show boundaries separating the three egg-abundance survey regions (eastern, western, and southwestern) described in the text.

Past fluctuations in abundance of the Pacific stock were accompanied by changes in spawning period (Itoh et al., 2009; Takahashi et al., 2008; Watanabe et al., 1996, 1997). The main spawning period occurred in April during the low biomass period and in February to March during the high biomass period (Nakai & Hattori, 1962; Watanabe et al., 1996). Changes in spawning period can affect the environment experienced by offspring (e.g., water temperature and food availability), which can lead to differences in growth (Itoh et al., 2011; Okunishi et al., 2012). In addition, early growth is important for early survival (Takahashi et al., 2008). Nevertheless, in Japanese sardine, it is not known how differences in the timing of spawning affect offspring growth rates.

Several studies have examined spawning patterns and early life history in the Japanese sardine. However, most of the important findings were obtained during the period of declining stock abundance when the stock was near its lowest level. New information collected during the recent period of increasing stock size is needed to understand the mechanisms driving population dynamics and recruitment fluctuations.

The present study aims to describe spatio-temporal changes in spawning pattern and early growth of Japanese sardine in the western North Pacific during the recent period of stock increase, and to clarify the effects of differences in spawning period on offspring growth. We first examined egg abundance in several regions off the Pacific coast and assessed hatch timing since 2004 to identify changes in spawning period in recent years (post-2010) when the stock has been recovering. We then examined the growth trajectory of individuals in the nursery by hatch-month cohort and identified the impact of spawning period changes on early growth.

## MATERIALS AND METHODS

### Sampling

Juvenile Japanese sardine (*S. melanostictus*) were collected by surface trawl survey in the western North Pacific in May and June during 2004 to 2018 (Table 1; Fig. 2). The survey method followed Takahashi et al. (2008); the net was towed at 3–4 kn and at depths of 0– 30 m for 30–60 min, and had a mouth opening size of 25 × 25 m with a 4-mm mesh at the cod end. The trawl surveys were conducted at 40–63 stations in each year. Catch per unit effort (CPUE) of juveniles at each station was calculated as the number of individuals collected per hour. Juveniles were frozen onboard the research vessel and standard length (SL) was measured in the laboratory to the nearest 0.01 mm by using digital calipers.

**Table 1.** Summary of survey and sampling data for juvenile Japanese sardine in the Kuroshio–Oyashio transitional region. CPUE: catch per unit effort (individuals h^−1^).

| Year | Survey dates | Station No. | Latitude range | Longitude range | Mean CPUE<br>(excluding zero<br>catch stations) | Relative frequency<br>of zero catch stations |
| --- | --- | --- | --- | --- | --- | --- |
| 2004 | 12 Mar–30 Mar | 50 | 35.02–39.55 | 142.57–164.00 | 11.50 | 0.84 |
| 2005 | 13 Mar–29 Mar | 45 | 35.01–39.22 | 143.01–162.01 | 389.14 | 0.69 |
| 2006 | 11 Mar–28 Mar | 42 | 34.58–39.17 | 143.58–163.01 | 128.50 | 0.90 |
| 2007 | 12 Mar–28 Mar | 43 | 35.01–39.08 | 143.04–162.02 | 355.70 | 0.67 |
| 2008 | 10 Mar–27 Mar | 40 | 35.04–39.02 | 142.59–159.58 | 117.00 | 0.95 |
| 2009 | 8 Mar–26 Mar | 49 | 35.08–39.17 | 142.59–160.04 | 199.34 | 0.49 |
| 2010 | 26 Mar–20 Jun | 57 | 35.47–39.41 | 143.16–160.29 | 3926.43 | 0.26 |
| 2011 | 25 Mar–16 Jun | 40 | 36.19–39.49 | 143.32–159.31 | 176.70 | 0.48 |
| 2012 | 24 Mar–18 Jun | 59 | 35.50–39.59 | 142.51–160.01 | 3361.21 | 0.44 |
| 2013 | 25 Mar–20 Jun | 59 | 34.52–41.07 | 143.31–160.06 | 2556.38 | 0.54 |
| 2014 | 24 Mar–18 Jun | 63 | 35.40–40.30 | 142.51–159.59 | 892.08 | 0.57 |
| 2015 | 23 Mar–18 Jun | 62 | 36.02–40.49 | 142.32–160.01 | 759.20 | 0.58 |
| 2016 | 25 Mar–19 Jun | 51 | 35.51–40.15 | 142.28–159.59 | 784.54 | 0.41 |
| 2017 | 24 Mar–17 Jun | 52 | 36.32–40.28 | 142.19–158.03 | 1040.20 | 0.58 |
| 2018 | 11 Mar–2 Jun | 43 | 36.00–39.01 | 143.00–158.58 | 13396.61 | 0.28 |

### Otolith microstructure analysis

In accordance with a previous study, sagittal otoliths were extracted from each specimen and processed for interpretation of daily growth increments (Takahashi et al., 2008). Otoliths were mounted on a glass plate with epoxy resin and polished using a lapping film. Increment widths (IW) were measured along the longest axis on the posterior side of the otolith. Age in days of each individual was assigned by adding 2 to the total number of otolith daily increments because the first daily increment forms 2–3 d after hatching in Japanese sardine (Hayashi et al., 1989). Daily growth increment profiles were established for each specimen by using an otolith measurement system (Ratoc System Engineering, Tokyo, Japan). Hatch date was estimated from the capture date and age of each fish, and somatic growth trajectory was back-calculated by using the biological intercept method (Campana, 1990). The back calculation of SL followed the methodology of a previous study (Takahashi et al., 2008). In Japanese sardine, the relationship between SL and otolith radius is linear after the juvenile stage (Takahashi et al., 2008), whereas the larval stage exhibits asymptotic growth (Watanabe & Kuroki, 1997). According to Watanabe & Kuroki (1997), larval total lengths converged to an infinite length of 29–32 mm. Therefore, we assumed a linear relationship for SLs over 30 mm, and only considered back-calculated SLs with means greater than 30 mm. Some of the specimens used in our study overlap with those of Furuichi et al. (2020a).

### Egg abundance data

To examine spawning periods of Japanese sardine, we used statistical reports on monthly egg production in coastal Pacific waters off Japan (Anonymous, 2004–2018). These reports are based on egg surveys conducted by a cooperative program involving three national research institutes in the Japan Fisheries Research and Education Agency and 18 prefectural fisheries research institutes in Japan since 1978 (Oozeki et al., 2007; Takasuka et al., 2008). In these surveys, conical or cylindrical–conical plankton nets with mouth ring diameters of 0.45 or 0.60 m and mesh sizes of 0.330 or 0.335 mm were towed vertically from ~150-m depth to the surface (Takasuka et al., 2017). We divided the reported data by location into three regions in the Pacific Ocean near Japan (eastern: >135°30′E; western: 131°30′–135°30′E; southwestern: <131°30′E, <31°30′N; Fig. 2) and examined monthly total egg abundance by year.

### Statistics

Measured IWs and back-calculated SLs were compared among hatch-month cohorts to detect whether differences among hatch-months affected growth trajectories. Differences among 10-day means of IW and back-calculated SL were tested by using Welch’s *t*-test or one-way analysis of variance (ANOVA, without assuming equal variances) with Scheffe’s test. For statistical analysis, we pooled data from all survey years.

## RESULTS

### Egg abundance

During 2004–2009, egg production of Japanese sardine in coastal Pacific waters off Japan was highest in the western region, which accounted for a large portion of total production (Fig. 3). Egg production in the western region tended to be high from January to March. From 2010 onwards, egg production increased in the eastern region in April and May; since 2013, pre-March egg production has also increased. Total egg production increased considerably after 2014, with the eastern region accounting for most of the increase. Spawning was rarely observed in the southwestern region throughout the study period.

**Fig. 3.**
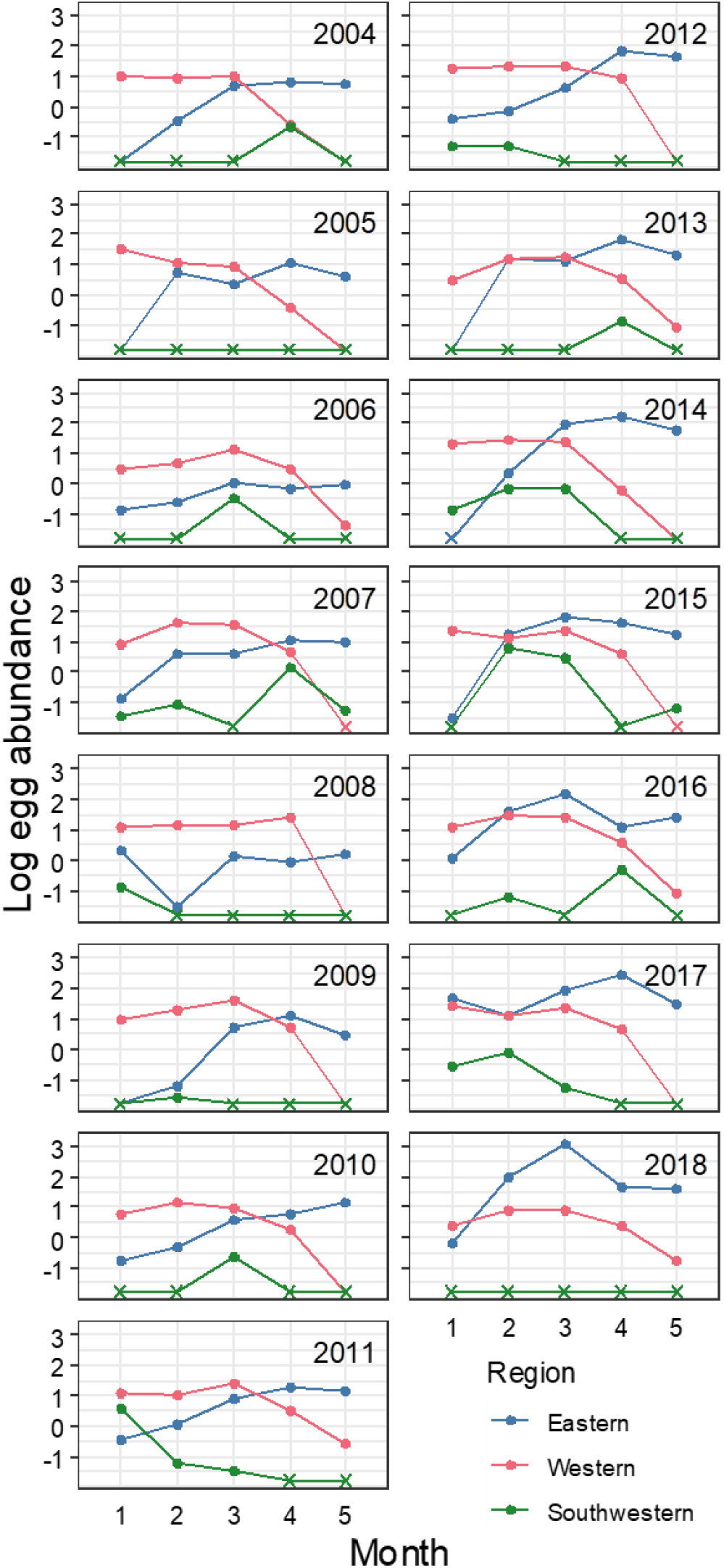
Abundance (×10^12^) of Japanese sardine eggs off the Japanese Pacific coast. Data are shown on a log10 scale (data from Anonymous, 2004–2018). Filled circles indicate abundances >0, and crosses indicate absence.

### Fish size, age, and hatch dates

From 2004 to 2012, almost all individuals were less than 70 d old and about 30–60 mm SL (Fig. 4). Hatch dates were concentrated from mid-March to April (Fig. 5). Until 2012, individuals that hatched in February or early March were rarely found. Since 2013, the number of individuals exceeding 60 mm SL and 70–120 d old has increased. Over time, more individuals have hatched in February and early March. The 2016 year-class was particularly large and old, exceeding 90 mm SL and with a modal age of around 100 d. In 2016 and 2018, few individuals hatched after April.

**Fig. 4.**
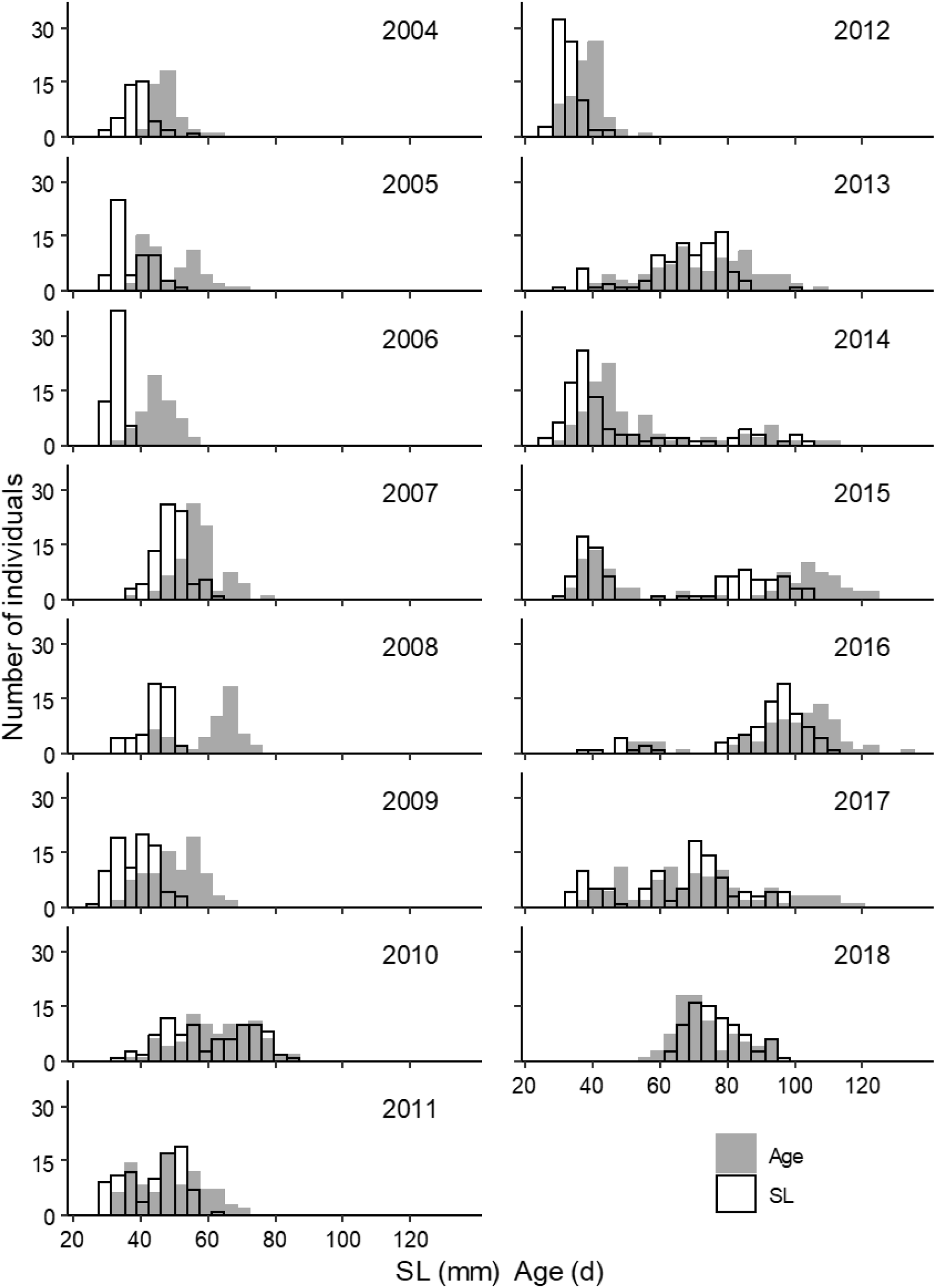
Frequency distributions of standard length (SL; open bars) and age (gray bars; in days) of juvenile Japanese sardine captured in the Kuroshio–Oyashio transitional region and used in our otolith analysis.

**Fig. 5.**
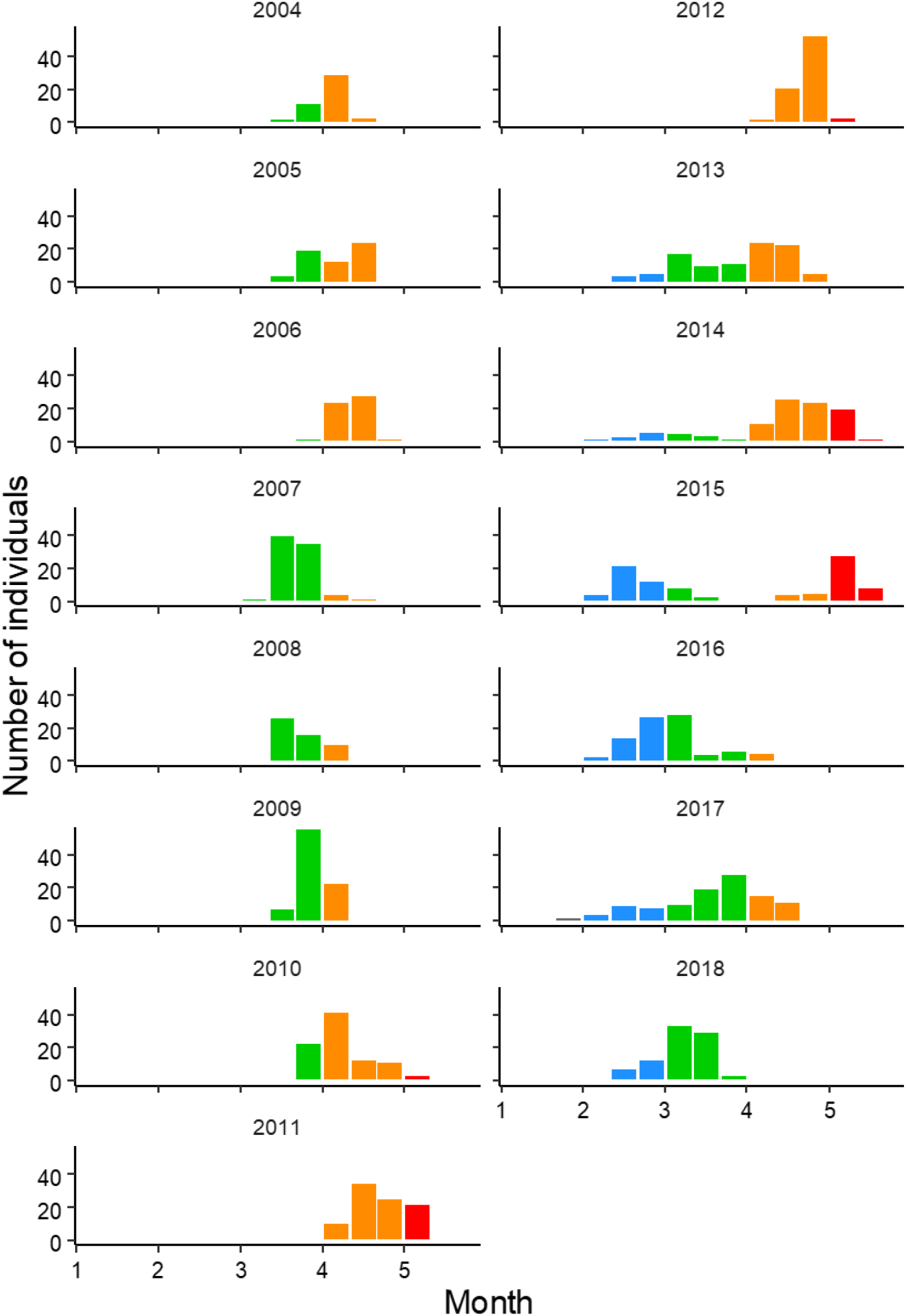
Frequency distribution of hatch dates of juvenile Japanese sardine captured in the Kuroshio–Oyashio transitional region and used in our otolith analysis. Colors indicate hatch months. Hatch date frequencies were calculated using 10-d bins.

### Growth trajectory of year-classes

Mean IWs were 3.4–5.5 μm at the age of 10 d and peaked at 7.8–12.1 μm at 40–60 d in many year-classes (Fig. 6a, Table S1). The age and magnitude of the peak differed among year-classes. Until 2012, when hatches predominantly occurred after mid-March, most year-classes reached peak IW by 40–50 d with values as low as than 10 μm. Especially in the 2011 and 2012 year-classes, in which many individuals hatched after mid-April, mean IWs increased rapidly up to an age of 30 d. On the other hand, since 2013, when some larvae hatched earlier, mean IWs peaked at 50–70 d, with maximum values exceeding 10 μm in most year-classes. In particular, mean IWs remained low up to an age of 20–30 d in the 2016 and 2018 year-classes, which had a higher proportion of individuals hatched before early March. At the age of 40 d, the difference in back-calculated SL between year-classes was about 5 mm, but at 50–60 d, the difference increased to more than 15 mm (Fig. 6b, Table S2).

**Fig. 6.**
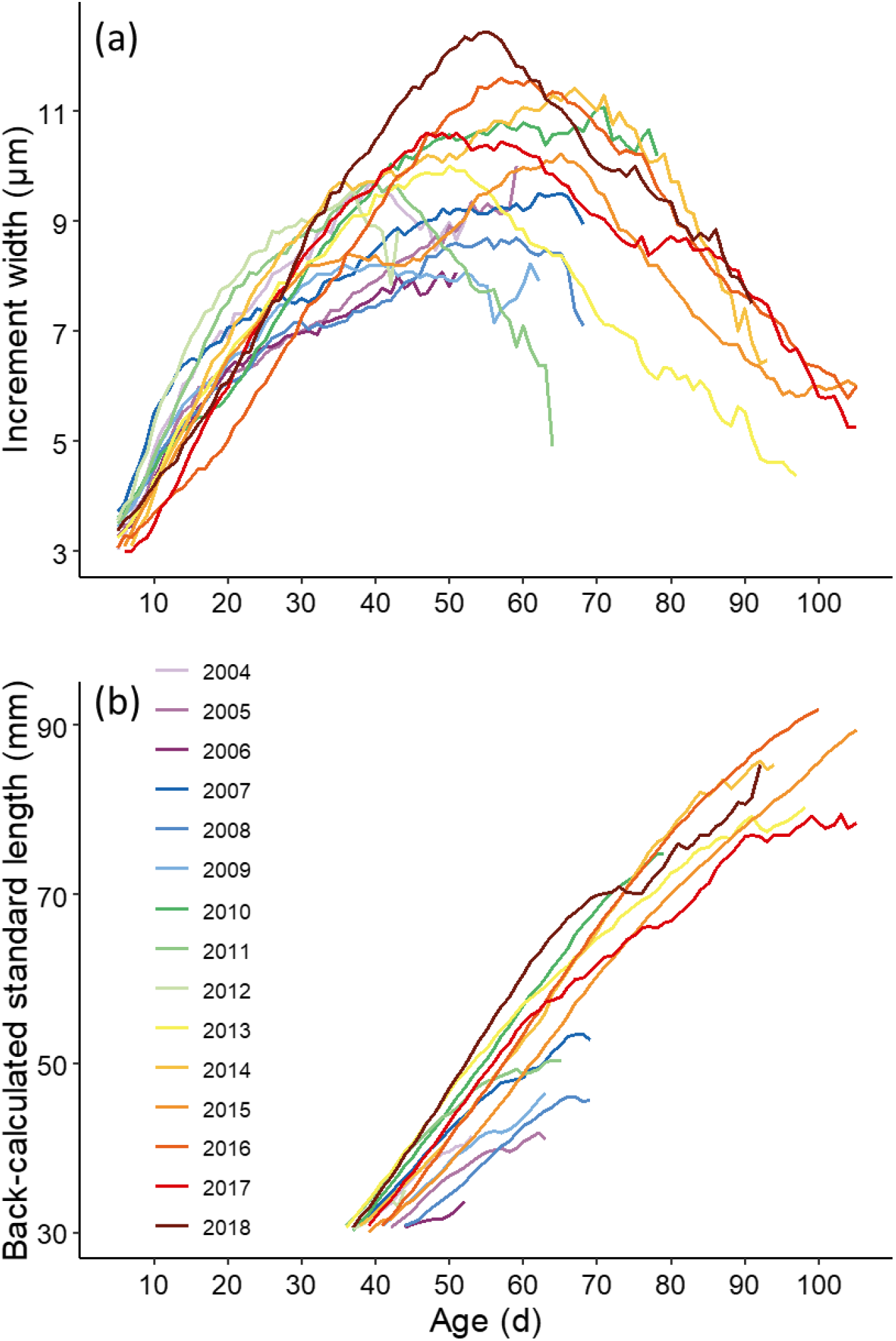
Relationships between age in days and (a) mean otolith increment width and (b) mean size-at-age (i.e., back-calculated standard length) of juvenile Japanese sardine in various year-classes. Means calculated from an *n* of less than 5 samples are not shown.

### Comparison of growth trajectory among hatch-month cohorts

Hatch-month cohorts were divided into 4 groups from February to May. In all cohorts, IW increased for individuals of age 10–30 d, with later hatch-month cohorts having the greater increase during this period (Fig. 7a, Table S3). After 40 d, the increase in IW stagnated and eventually declined, with the decline happening first in later hatch-month cohorts. Earlier hatch-month cohorts, especially the February-hatched cohort, experienced a modest increase in IW. However, the February-hatched cohort had a longer period of IW increase and exceeded March- and April- hatched cohorts at ages of around 60–70 d (ANOVA, 10 d: *F* = 92.61, *P* < 0.01; 20 d: *F* = 174.73, *P* < 0.01; 30 d: *F* = 96.08, *P* < 0.01; 40 d: *F* = 21.77, *P* < 0.01; 50 d: *F* = 8.15, *P* < 0.01; 60 d: *F* = 13.26, *P* < 0.01; 70 d: *F* = 17.63, *P* < 0.01; *t*-test, 80 d: *P* < 0.01; 90 d: *P* = 0.30; 100 d: *P* = 0.70).

**Fig. 7.**
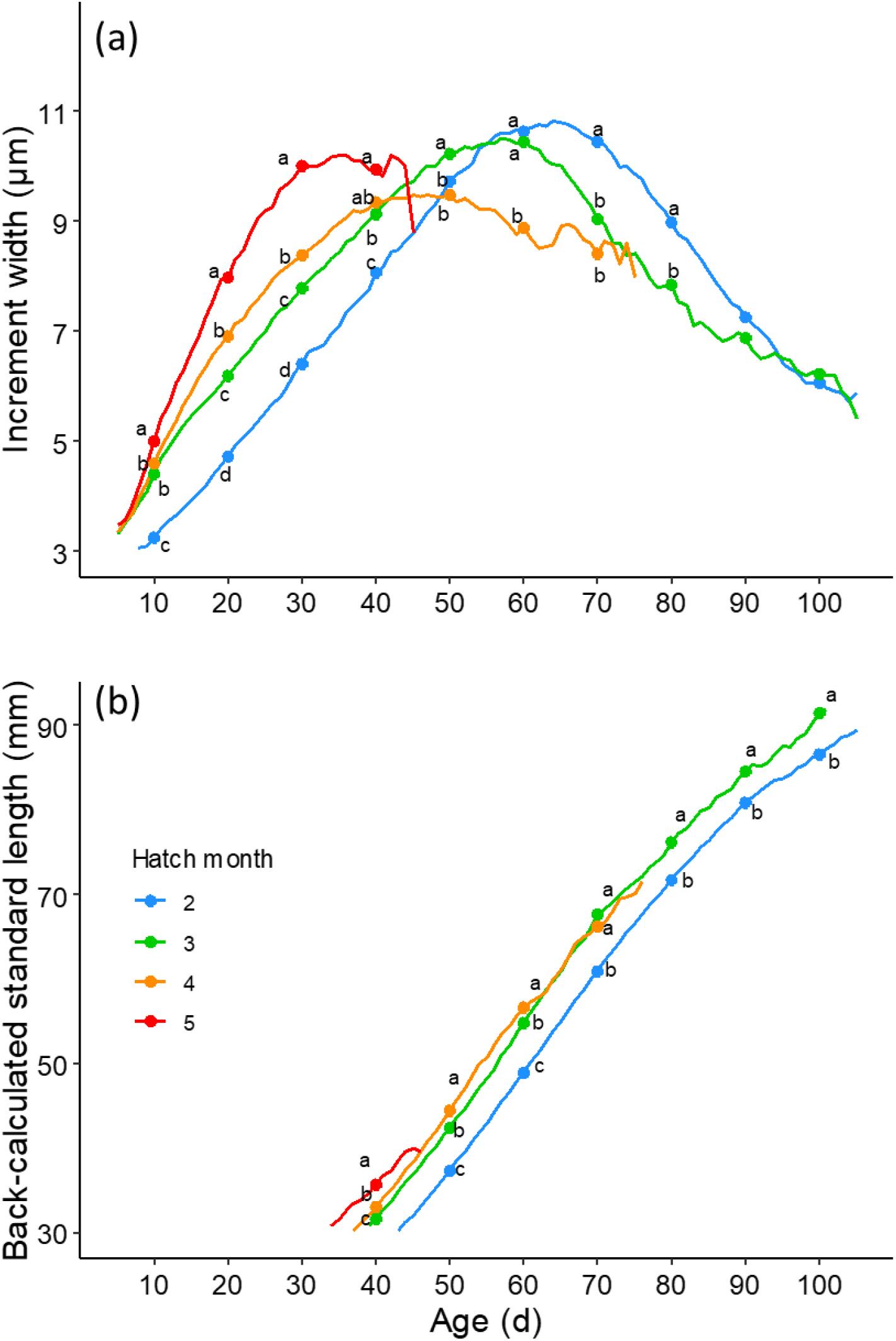
Relationships between age in days and (a) mean otolith increment widths and (b) mean size-at-age (i.e., back-calculated standard length) of juvenile Japanese sardine in various hatch-month cohorts. Means calculated from an *n* of less than 5 samples are not shown. Each timeseries has been annotated with letters at 10-d intervals; data points annotated with different letters are significantly different at *P* < 0.05.

Back-calculated SLs of later-hatched cohorts were larger than those of earlier-hatched cohorts before the age of 40–60 d (Fig. 7b, Table S4). At 70 d, March- and April-hatched cohorts were comparable, but the February-hatched cohort was consistently smaller than the March-hatched cohort (ANOVA, 40 d: *F* = 31.46, *P* < 0.01; 50 d: *F* = 67.28, *P* < 0.01; 60 d: *F* = 41.50, *P* < 0.01; 70 d: *F* = 28.54, *P* < 0.01; *t*-test, 80 d: *P* < 0.01; 90 d: *P* < 0.05; 100 d: *P* < 0.01).

## DISCUSSION

In the eastern region, increases in egg production coincided with the recent period of recruitment increase; in addition, the spawning period shifted earlier in the year. This suggests that the eastern region is an important spawning ground for recruitment processes and that earlier-hatched cohorts in particular have contributed to the recent increase in recruitment. Early spawning (i.e., in February) was also consistently observed in the western region during the study period; however, no juveniles appear to have hatched in February prior to 2012. Therefore, it seems that individuals spawned in February in the western region rarely survived in the Kuroshio–Oyashio transitional region. Additionally, during our study period, spawning was seldom observed in the southwestern region, which was a major spawning ground during the last high-biomass period (Watanabe et al., 1996). Since the egg production peak of Japanese sardine is related to sea surface temperature (Furuichi et al., 2020b), one possible explanation for the lack of spawning in the southwestern region is an increase in water temperature due to climate change. Similar increases in water temperature have caused a northward shift since the 2000s in the spawning grounds of chub mackerel (*Scomber japonicus*), which spawns in the North Pacific off the coast of Japan (Kanamori et al., 2019). The expansion of Japanese sardine spawning to the offshore waters of the southwestern region is thought to have led to extremely high recruitment during the last high-biomass period (Watanabe et al., 1996). If current conditions inhibit the establishment of a southwestern spawning ground, future recruitment trends may differ from those of the past.

Hatch dates of juveniles caught in the study area were generally consistent with spawning periods in the eastern region. This suggests that the eastern region is an important spawning ground for juveniles in the Kuroshio–Oyashio transitional region and plays a key role in the recruitment processes of this stock. However, in some years, hatch dates and spawning periods did not coincide, as in 2015. This may be due to biological factors such as failure to survive as a result of mismatch, or could have been caused by sampling bias. A study using a transport model of the Pacific stock of Japanese sardine suggested that the dispersal range of larvae spawned in the eastern region can vary greatly over a period of tens of days (Itoh & Kimura, 2007), and it is possible that our survey did not adequately capture the dynamics of the cohort that provided the main source of recruitment in 2015.

In this study, cohorts that hatched earlier experienced slower early growth than did cohorts that hatched later. The relationship between early growth rate of Japanese sardine and water temperature follows a dome-shaped curve, with an optimum temperature range of approximately 16–18°C (Takahashi et al., 2009; Takasuka et al., 2007). Surface water temperatures in the Kuroshio region rise from around 18°C in February–March to about 20–25°C from April to June. Drifter measurements have shown that eggs and larvae spawned in February–March and transported toward the eastern Kuroshio region do not experience large increases in water temperature (Itoh & Kimura, 2007). In contrast, eggs and larvae spawned in April–May would experience continuously higher water temperatures than those spawned in February–March. This means that eggs and larvae spawned in February–March may experience suitable water temperatures for growth for a relatively longer period as they are advected to their nursery grounds. On the other hand, algal blooms in the Kuroshio Extension peak in April–May (Shiozaki et al., 2014), and copepods, the primary food of larval Japanese sardine, are most abundant during this period (Nakata et al., 2004). The comparatively low prey availability in February–March probably accounts for the slow growth of earlier-hatched cohorts because of the positive relationship between food density and growth rate at appropriate water temperatures (Takahashi et al., 2009).

The relatively slow growth of recent year-classes during early life seems to reflect the preponderance of earlier-hatched cohorts. On the other hand, both IW and back-calculated SL after 50 d in post-2013 year-classes tended to be larger than in year-classes prior to 2012 (except for 2010). Furthermore, back-calculated SLs at ages of 50–60 d exceeded those of samples collected during the low biomass period (1996–2003) (Takahashi et al., 2008). Thus, the growth rates of Japanese sardine during the recent period of stock increase seem to follow a pattern of slow growth during the larval stage followed by rapid growth during the juvenile stage.

Our results suggest that early-spawned individuals are important in increasing recruitment despite their low early growth rates. A study on the Pacific stock of Japanese sardine during the low biomass period reported a positive correlation between growth after metamorphosis (50–60 d of age) and recruitment (Takahashi et al., 2008). However, Furuichi et al. (2020a) indicated that the growth–recruitment relationship varies dynamically over time; the positive effect of growth rate on recruitment success has diminished since the late 2000s and the baseline recruitment level has increased. Some species, such as walleye pollock (*Gadus chalcogramma*) and Norwegian herring (*Clupea harengus*), had higher recruitment in low-growth cohorts than in high-growth cohorts (Nishimura et al., 2007; Slotte et al., 2019). Similarly, low-growth Japanese sardine (i.e., early-spawned individuals) may survive at higher rates than high-growth cohorts.

Differences in the timing of spawning can lead to differences in the environment that offspring experience during their early life stages. In turn, differences in environmental conditions can affect offspring survival through changes in metabolic rates and the frequency of predator encounters (Durant et al., 2007; Houde, 1989). Various taxa are thought to prey on Japanese sardine, including carnivorous microzooplankton, micronekton, large pelagic fishes, marine mammals, and seabirds (Sugisaki, 1996). In the Kuroshio region, for instance, clupeoids including Japanese sardine are important prey species for Pacific bluefin tuna (*Thunnus orientalis*) (Shimose et al., 2013). Juvenile bluefin that remain in the coastal waters of the western region in summer and autumn begin to migrate to the eastern region from December to February, and are increasingly found in the Kuroshio–Oyashio transitional region from March onward (Fujioka et al., 2018). Early spawning of Japanese sardine may reduce the frequency of encounters with predators during early life. Furthermore, once the bluefin begin to enter the Kuroshio– Oyashio transitional region, early-spawned individuals may be more able to escape from predators (despite their lower growth rate) due to their more advanced growth state. For these reasons, earlier-hatched cohorts may experience less predation-based mortality. In addition, water temperatures could affect survival rates through changes in metabolism. At high water temperatures (which raise metabolic rates), individuals with inadequate food supply are at increased risk of starvation. Conversely, whereas growth is suppressed at low water temperatures, reduced metabolic rates increase the chances of survival during periods of low food supply (Garrido et al., 2016; Houde, 1989). Early-spawned Japanese sardine experience relatively low water temperatures, which may help them maintain high survival rates despite the limited food availability caused by missing the peak season for plankton blooms.

In conclusion, our results suggest that spawning grounds in the eastern part of the waters off Japan (herein, the “eastern region”) play a key role in the recruitment of the Pacific stock of Japanese sardine, and that early-hatched cohorts are contributing especially strongly to recent increases in recruitment. These early-hatched larvae and juveniles have unexpectedly lower growth rates during their early life stages. Their recruitment success is likely to be supported by factors other than growth rate, such as a reduced frequency of predator encounters due to earlier spawning. Our study underlines the importance of spawning patterns on influencing recruitment processes. Further research to identify reproductive traits associated with changes in stock size (and consequent changes in recruitment processes) will help refine predictions of population dynamics and reveal the mechanisms underlying recruitment fluctuations more generally.

## Acknowledgements

The authors are grateful to the officers and crews of training vessel *Hokuho-Maru* and research vessel *Soyo-Ma*ru, as well as to participating researchers for their support in the field. We also thank Mika Suhara and Taishi Okumura for their help during the otolith microstructure analysis. Funding for this study was provided by the Japan Fisheries Research and Education Agency and the Fisheries Agency of Japan.

## REFERENCES

Anonymous (2004–2018). Annual Meeting Report of Egg and Larval Survey of Small Pelagic Fish off the Pacific Coast of Japan, No. 24–38. (in Japanese). National Research Institute of Fisheries Science, Japan Fisheries Research and Education Agency of Japan.

Campana, S. E. (1990). How reliable are growth back-calculations based on otoliths? Canadian Journal of Fisheries and Aquatic Sciences, 47(11), 2219–2227. https://doi.org/10.1139/f90-246

Durant, J. M., Hjermann, D., Ottersen, G., & Stenseth, N. C. (2007). Climate and the match or mismatch between predator requirements and resource availability. Climate Research, 33(3), 271–283. https://doi.org/10.3354/cr033271

Fujioka, K., Fukuda, H., Furukawa, S., Tei, Y., Okamoto, S., & Ohshimo, S. (2018). Habitat use and movement patterns of small (age-0) juvenile Pacific bluefin tuna (*Thunnus orientalis*) relative to the Kuroshio. Fisheries Oceanography, 27(3), 185–198. https://doi.org/10.1111/fog.12244

Furuichi, S., Niino, Y., Kamimura, Y., & Yukami, R. (2020). Time-varying relationships between early growth rate and recruitment in Japanese sardine. Fisheries Research, 232, 105723. https://doi.org/10.1016/j.fishres.2020.105723

Furuichi, S., Yasuda, T., Kurota, H., Yoda, M., Suzuki, K., Takahashi, M., & Fukuwaka, M. A. (2020). Disentangling the effects of climate and density-dependent factors on spatiotemporal dynamics of Japanese sardine spawning. Marine Ecology Progress Series, 633, 157–168. https://doi.org/10.3354/meps13169

Furuichi, S., Yukami, R., Kamimura, Y., Hayashi, A., Isu, S., and Watanabe, R. (2020c). Stock assessment and evaluation for the Pacific stock of sardine (fiscal year 2019). In Marine fisheries stock assessment and evaluation for Japanese waters (fiscal year 2019/20). (in Japanese)

Garrido, S., Cristóvão, A., Caldeira, C., Ben-Hamadou, R., Baylina, N., Batista, H., Saiz, E., Peck, M.A., Ré, P., & Santos, A. M. P. (2016). Effect of temperature on the growth, survival, development and foraging behaviour of *Sardina pilchardus* larvae. Marine Ecology Progress Series, 559, 131–145. https://doi.org/10.3354/meps11881

Hayashi, A., Yamashita, Y., Kawaguchi, K., & Ishii, T. (1989). Rearing Method and Daily Otolith Ring of Japanese Sardine Larvae. Nippon Suisan Gakkaishi, 55(6), 997–1000. https://doi.org/10.2331/suisan.55.997

Houde, E. D. (1989). Comparative growth, mortality, and energetics of marine fish larvae: temperature and implied latitudinal effects. Fishery Bulletin, 87(3), 471–495.

Itoh, S., & Kimura, S. (2007). Transport and survival of larvae of pelagic fishes in Kuroshio system region estimated with Lagrangian drifters. Fisheries Science, 73(6), 1295–1308. https://doi.org/10.1111/j.1444-2906.2007.01468.x

Itoh, S., Saruwatari, T., Nishikawa, H., Yasuda, I., Komatsu, K., Tsuda, A., Setou, T., and Shimizu, M. (2011). Environmental variability and growth histories of larval Japanese sardine (*Sardinops melanostictus*) and Japanese anchovy (*Engraulis japonicus*) near the frontal area of the Kuroshio. Fisheries Oceanography, 20(2), 114–124. https://doi.org/10.1111/j.1365-2419.2011.00572.x

Itoh, S., Yasuda, I., Nishikawa, H., Sasaki, H., & Sasai, Y. (2009). Transport and environmental temperature variability of eggs and larvae of the Japanese anchovy (*Engraulis japonicus*) and Japanese sardine (*Sardinops melanostictus*) in the western North Pacific estimated via numerical particle-tracking experiments. Fisheries Oceanography, 18(2), 118–133. https://doi.org/10.1111/j.1365-2419.2009.00501.x

Kanamori, Y., Takasuka, A., Nishijima, S., & Okamura, H. (2019). Climate change shifts the spawning ground northward and extends the spawning period of chub mackerel in the western North Pacific. Marine Ecology Progress Series, 624, 155–166. https://doi.org/10.3354/meps13037

Lluch-Belda, D., Crawford, R. J. M., Kawasaki, T., MacCall, A. D., Parrish, R. H., Schwartzlose, R. A., & Smith, P. E. (1989). World-wide fluctuations of sardine and anchovy stocks: The regime problem. South African Journal of Marine Science, 8(1), 195–205. https://doi.org/10.2989/02577618909504561

Nakai, Z., & Hattori, S. (1962). Quantitative distribution of eggs and larvae of the Japanese sardine by year, 1949 through 1951. Tokai Reg. Fish. Res. Lab., 9, 23–60.

Nakata, K., Itoh, H., Ichikawa, T., & Sasaki, K. (2004). Seasonal changes in the reproduction of three oncaeid copepods in the surface layer of the Kuroshio Extension. Fisheries Oceanography, Supplement, 13(1), 21–33. https://doi.org/10.1111/j.1365-2419.2004.00316.x

Nishimura, A., Hamatsu, T., Shida, O., Mihara, I., & Mutoh, T. (2007). Interannual variability in hatching period and early growth of juvenile walleye pollock, *Theragra chalcogramma*, in the Pacific coastal area of Hokkaido. Fisheries Oceanography, 16(3), 229–239. https://doi.org/10.1111/j.1365-2419.2007.00428.x

Okunishi, T., Ito, S. I., Ambe, D., Takasuka, A., Kameda, T., Tadokoro, K., Setou, T., Komatsu, K., Kawabata, A., Kubota, H., Ichikawa, T., Sugisaki, H., Hashioka, T.,Yamanaka, Y., Yoshie, N., & Watanabe, T. (2012). A modeling approach to evaluate growth and movement for recruitment success of Japanese sardine (*Sardinops melanostictus*) in the western Pacific. Fisheries Oceanography, 21(1), 44–57. https://doi.org/10.1111/j.1365-2419.2011.00608.x

Oozeki, Y., Takasuka, A., Kubota, H., & Barange, M. (2007). Characterizing spawning habitats of Japanese sardine (*Sardinops melanostictus*), Japanese anchovy (*Engraulis japonicus*), and pacific round herring (*Etrumeus teres*) in the Northwestern Pacific. California Cooperative Oceanic Fisheries Investigations Reports, 48, 191–203.

Sakamoto, T., Komatsu, K., Shirai, K., Higuchi, T., Ishimura, T., Setou, T., Kamimura, Y., Watanabe, C., & Kawabata, A. (2019). Combining microvolume isotope analysis and numerical simulation to reproduce fish migration history. Methods in Ecology and Evolution, 10(1), 59–69. https://doi.org/10.1111/2041-210X.13098

Schwartzlose, R. A., Alheit, J., Bakun, A., Baumgartner, T. R., Cloete, R., Crawford, R. J. M., Fletcher, W. J., Green-Ruiz, Y., Hagen, E., Kawasaki, T., Lluch-Belda, D., Lluch-Cota, S. E., MacCall, A. D., Matsuura, Y., Nevárez-Martínez, M. O., Parrish, R. H., Roy, C., Serra, R., Shust, K. V., Ward, M. N. & Zuzunaga, J. Z. (1999). WORLDWIDE LARGE-SCALE FLUCTUATIONS OF SARDINE AND ANCHOVY POPULATIONS In 1993, the Scientific Committee on Oceanic Research (SCOR) established Working Group 98 (WG 98) to investigate “Worldwide Large-scale Fluctuations of Sardine and Anchovy Popul.” (October 1998), 289–347.

Secor, D. H. (2007). The year-class phenomenon and the storage effect in marine fishes. Journal of Sea Research, 57(2-3 SPEC. ISS.), 91–103. https://doi.org/10.1016/j.seares.2006.09.004

Shimose, T., Watanabe, H., Tanabe, T., & Kubodera, T. (2013). Ontogenetic diet shift of age-0 year Pacific bluefin tuna *Thunnus orientalis*. Journal of Fish Biology, 82(1), 263–276. https://doi.org/10.1111/j.1095-8649.2012.03483.x

Shiozaki, T., Ito, S. I., Takahashi, K., Saito, H., Nagata, T., & Furuya, K. (2014). Regional variability of factors controlling the onset timing and magnitude of spring algal blooms in the northwestern North Pacific. Journal of Geophysical Research: Oceans, 119(1), 253–265. https://doi.org/10.1002/2013JC009187

Slotte, A., Husebø, Å., Berg, F., Stenevik, E. K., Folkvord, A., Fossum, P., Mosegaard, H., Vikebø, F., & Nash, R. D. M. (2019). Earlier hatching and slower growth: A key to survival in the early life history of Norwegian spring spawning herring. Marine Ecology Progress Series, 2019(617–618), 25–39. https://doi.org/10.3354/meps12682

Sugisaki, H. (1996). Distribution of larval and juvenile Japanese sardine (*Sardinops melanostictus*) in the western North Pacific and its relevance to predation on these stages. In Y. Watanabe, Y. Yamashita, & Y. Oozeki (Eds.), Survival strategies in early life stage of marine resources (pp. 261–270). Balkema.

Takahashi, M., Nishida, H., Yatsu, A., & Watanabe, Y. (2008). Year-class strength and growth rates after metamorphosis of Japanese sardine (*Sardinops melanostictus*) in the western North Pacific Ocean during 1996-2003. Canadian Journal of Fisheries and Aquatic Sciences, 65(7), 1425–1434. https://doi.org/10.1139/F08-063

Takahashi, M., Watanabe, Y., Yatsu, A., & Nishida, H. (2009). Contrasting responses in larval and juvenile growth to a climate-ocean regime shift between anchovy and sardine. Canadian Journal of Fisheries and Aquatic Sciences, 66(6), 972–982. https://doi.org/10.1139/F09-051

Takasuka, A., Kubota, H., & Oozeki, Y. (2008). Spawning overlap of anchovy and sardine in the western North Pacific. Marine Ecology Progress Series, 366, 231–244. https://doi.org/10.3354/meps07514

Takasuka, A., Oozeki, Y., & Aoki, I. (2007). Optimal growth temperature hypothesis: Why do anchovy flourish and sardine collapse or vice versa under the same ocean regime? Canadian Journal of Fisheries and Aquatic Sciences, 64(5), 768–776. https://doi.org/10.1139/F07-052

Takasuka, A., Tadokoro, K., Okazaki, Y., Ichikawa, T., Sugisaki, H., Kuroda, H., & Oozeki, Y. (2017). In situ filtering rate variability in egg and larval surveys off the Pacific coast of Japan: Do plankton nets clog or over-filter in the sea? Deep-Sea Research Part I: Oceanographic Research Papers, 120, 132–137. https://doi.org/10.1016/j.dsr.2016.12.017

Watanabe, Y., & Kuroki, T. (1997). Asymptotic growth trajectories of larval sardine (*Sardinops melanostictus*) in the coastal waters off western Japan. Marine Biology, 127(3), 369–378. https://doi.org/10.1007/s002270050023

Watanabe, Y., Zenitani, H., & Kimura, R. (1995). Population decline off the Japanese sardine *Sardinops melanostictus* owing to recruitment failures. Canadian Journal of Fisheries and Aquatic Sciences, 52(8), 1609–1616. https://doi.org/10.1139/f95-154

Watanabe, Y., Zenitani, H., & Kimura, R. (1996). Offshore expansion of spawning of the Japanese sardine, *Sardinops melanostictus*, and its implication for egg and larval survival. Canadian Journal of Fisheries and Aquatic Sciences, 53(1), 55–61. https://doi.org/10.1139/f95-153

Watanabe, Y., Zenitani, H., & Kimura, R. (1997). Variations in spawning ground area and egg density of the Japanese sardine in Pacific coastal and oceanic waters. Fisheries Oceanography, 6(1), 35–40. https://doi.org/10.1046/j.1365-2419.1997.00024.x

Yasuda, I., Sugisaki, H., Watanabe, Y., Minobe, S. S., & Oozeki, Y. (1999). Interdecadal variations in Japanese sardine and ocean/climate. Fisheries Oceanography, 8(1), 18–24. https://doi.org/10.1046/j.1365-2419.1999.00089.x

Yatsu, A. (2019). Review of population dynamics and management of small pelagic fishes around the Japanese Archipelago. Fisheries Science, 85(4), 611–639. https://doi.org/10.1007/s12562-019-01305-3

