## Supplemental Table for "Spatio-temporal spawning patterns and early growth of Japanese sardine in the western North Pacific during the recent stock increase"

Table S1. Mean otolith increment widths for various year-classes of juvenile Japanese sardine in the Kuroshio–Oyashio transitional region

| | Increment width (μm) by age in days | | | | | | | | | | | | | | | | | | | |
|---|---|---|---|---|---|---|---|---|---|---|---|---|---|---|---|---|---|---|---|---|
| | 10 d | | 20 d | | 30 d | | 40 d | | 50 d | | 60 d | | 70 d | | 80 d | | 90 d | | 100 d | |
| Year | Mean ± SD | *n* | Mean ± SD | *n* | Mean ± SD | *n* | Mean ± SD | *n* | Mean ± SD | *n* | Mean ± SD | *n* | Mean ± SD | *n* | Mean ± SD | *n* | Mean ± SD | *n* | Mean ± SD | *n* |
| 2004 | 4.8 ± 0.9 | 43 | 6.8 ± 1.2 | 43 | 8.4 ± 1.5 | 43 | 9.7 ± 1.7 | 41 | 8.9 ± 1.4 | 9 | 11.3 | 1 | | | | | | | | |
| 2005 | 4.4 ± 0.9 | 57 | 6.2 ± 1.0 | 57 | 6.9 ± 1.1 | 57 | 7.9 ± 1.3 | 52 | 8.9 ± 1.4 | 25 | 10.0 ± 1.1 | 4 | | | | | | | | |
| 2006 | 4.2 ± 0.8 | 54 | 6.3 ± 1.2 | 54 | 6.9 ± 0.9 | 54 | 7.6 ± 0.9 | 47 | 7.8 ± 1.5 | 9 | | | | | | | | | | |
| 2007 | 5.5 ± 1.1 | 80 | 7.0 ± 1.2 | 80 | 7.6 ± 1.2 | 80 | 8.4 ± 1.3 | 79 | 9.1 ± 1.2 | 71 | 9.1 ± 1.2 | 19 | 7.9 ± 0.1 | 2 | | | | | | |
| 2008 | 4.5 ± 1.1 | 52 | 6.0 ± 1.2 | 52 | 7.1 ± 1.4 | 52 | 7.4 ± 1.6 | 52 | 8.5 ± 1.3 | 42 | 8.5 ± 1.4 | 37 | 7.4 ± 0.4 | 2 | | | | | | |
| 2009 | 4.6 ± 0.9 | 85 | 6.4 ± 1.3 | 85 | 7.8 ± 1.5 | 85 | 8.1 ± 1.7 | 73 | 7.9 ± 1.2 | 43 | 7.8 ± 1.5 | 9 | | | | | | | | |
| 2010 | 4.5 ± 0.8 | 88 | 5.8 ± 1.1 | 88 | 7.9 ± 1.3 | 88 | 9.7 ± 1.3 | 86 | 10.6 ± 1.2 | 76 | 10.8 ± 1.2 | 50 | 11.0 ± 1.0 | 24 | 10.8 ± 0.7 | 3 | | | | |
| 2011 | 4.7 ± 0.9 | 90 | 7.5 ± 1.3 | 90 | 8.7 ± 1.5 | 90 | 9.7 ± 1.3 | 66 | 8.4 ± 2.1 | 39 | 7.1 ± 3.3 | 13 | 3.8 | 1 | | | | | | |
| 2012 | 5.3 ± 1.2 | 75 | 7.8 ± 1.3 | 75 | 9.0 ± 1.6 | 72 | 8.9 ± 2.6 | 18 | 7.7 | 1 | | | | | | | | | | |
| 2013 | 4.2 ± 1.0 | 94 | 6.5 ± 1.2 | 94 | 8.0 ± 1.4 | 94 | 9.4 ± 1.5 | 93 | 10.0 ± 1.8 | 85 | 8.8 ± 2.3 | 75 | 7.2 ± 2.2 | 53 | 6.3 ± 1.7 | 32 | 5.5 ± 1.6 | 11 | 4.5 | 1 |
| 2014 | 4.0 ± 1.1 | 97 | 6.8 ± 1.6 | 97 | 8.8 ± 1.4 | 96 | 9.7 ± 1.7 | 71 | 10.2 ± 2.1 | 34 | 11.1 ± 2.0 | 20 | 11.0 ± 1.8 | 17 | 9.7 ± 2.1 | 14 | 7.4 ± 1.9 | 9 | 4.1 ± 0.9 | 2 |
| 2015 | 4.1 ± 1.1 | 90 | 6.4 ± 1.9 | 90 | 8.0 ± 2.3 | 90 | 8.3 ± 1.6 | 71 | 8.7 ± 1.9 | 52 | 9.9 ± 2.0 | 48 | 9.5 ± 1.9 | 46 | 7.8 ± 1.6 | 44 | 6.5 ± 1.4 | 43 | 5.9 ± 1.4 | 31 |
| 2016 | 3.6 ± 0.8 | 85 | 5.0 ± 1.1 | 85 | 7.2 ± 1.8 | 85 | 9.1 ± 2.0 | 85 | 10.9 ± 1.8 | 85 | 11.5 ± 1.6 | 78 | 10.7 ± 1.8 | 75 | 9.2 ± 2.1 | 75 | 7.6 ± 1.9 | 65 | 6.3 ± 1.5 | 42 |
| 2017 | 3.4 ± 1.0 | 105 | 5.9 ± 1.9 | 105 | 8.3 ± 2.4 | 105 | 9.6 ± 2.4 | 101 | 10.5 ± 2.6 | 83 | 10.3 ± 2.5 | 74 | 9.1 ± 2.5 | 52 | 8.7 ± 2.2 | 26 | 7.7 ± 2.5 | 20 | 5.8 ± 2.4 | 12 |
| 2018 | 4.2 ± 1.1 | 83 | 6.0 ± 1.4 | 83 | 8.4 ± 2.1 | 83 | 10.5 ± 2.4 | 83 | 12.1 ± 2.4 | 83 | 11.8 ± 1.9 | 81 | 10.1 ± 1.9 | 43 | 9.3 ± 1.6 | 21 | 7.7 ± 1.0 | 6 | | |

Table S2. Mean back-calculated standard lengths for various year-classes of juvenile Japanese sardine in the Kuroshio–Oyashio transitional region

| | Back-calculated standard length (mm) by age in days | | | | | | | | | | | | | |
|---|---|---|---|---|---|---|---|---|---|---|---|---|---|---|
| | 40 d | | 50 d | | 60 d | | 70 d | | 80 d | | 90 d | | 100 d | |
| Year | Mean ± SD | *n* | Mean ± SD | *n* | Mean ± SD | *n* | Mean ± SD | *n* | Mean ± SD | *n* | Mean ± SD | *n* | Mean ± SD | *n* |
| 2004 | 31.7 ± 3.2 | 42 | 39.5 ± 4.1 | 13 | 47.1 ± 7.8 | 2 | | | | | | | | |
| 2005 | | | 36.7 ± 4.0 | 39 | 40.8 ± 5.1 | 7 | | | | | | | | |
| 2006 | | | 31.9 ± 1.9 | 9 | | | | | | | | | | |
| 2007 | 32.8 ± 3.4 | 79 | 42.1 ± 3.9 | 74 | 48.3 ± 4.1 | 22 | 51.1 ± 2.5 | 2 | | | | | | |
| 2008 | | | 34.5 ± 2.7 | 43 | 42.5 ± 2.9 | 39 | 43.3 ± 1.9 | 2 | | | | | | |
| 2009 | | | 38.3 ± 3.3 | 47 | 43.5 ± 2.5 | 11 | | | | | | | | |
| 2010 | 33.4 ± 3.1 | 87 | 44.7 ± 3.6 | 77 | 56.9 ± 4.7 | 53 | 68.2 ± 4.6 | 27 | 76.6 ± 5.5 | 4 | | | | |
| 2011 | 34.6 ± 3.6 | 68 | 44.1 ± 4.1 | 47 | 48.6 ± 2.8 | 15 | 47.8 | 1 | | | | | | |
| 2012 | 32.0 ± 3.7 | 28 | 34.8 | 1 | | | | | | | | | | |
| 2013 | 34.9 ± 4.8 | 93 | 46.6 ± 5.8 | 86 | 57.1 ± 5.8 | 79 | 64.8 ± 5.1 | 54 | 72.5 ± 4.5 | 34 | 78.7 ± 7.5 | 11 | 84.2 ± 0.9 | 2 |
| 2014 | 32.3 ± 3.8 | 74 | 41.2 ± 6.1 | 35 | 52.8 ± 8.6 | 21 | 65.7 ± 9.0 | 17 | 77.7 ± 9.9 | 14 | 84.1 ± 9.4 | 10 | 82.8 ± 3.6 | 3 |
| 2015 | 30.7 ± 4.8 | 74 | 38.1 ± 4.3 | 52 | 48.7 ± 4.7 | 48 | 60.3 ± 4.7 | 46 | 70.1 ± 4.8 | 44 | 78.1 ± 5.2 | 43 | 85.7 ± 5.5 | 32 |
| 2016 | | | 41.1 ± 6.5 | 85 | 53.4 ± 7.0 | 79 | 66.1 ± 6.8 | 75 | 77.2 ± 6.2 | 75 | 85.6 ± 5.8 | 66 | 92 ± 5.4 | 45 |
| 2017 | 31.7 ± 6.3 | 102 | 43.0 ± 8.8 | 84 | 54.7 ± 10.2 | 77 | 61.7 ± 9.2 | 55 | 66.9 ± 9.9 | 29 | 76.7 ± 12.7 | 21 | 78.5 ± 14.9 | 13 |
| 2018 | 34.1 ± 5.8 | 83 | 47.0 ± 8.1 | 83 | 60.5 ± 9.6 | 81 | 69.9 ± 10.8 | 49 | 74.7 ± 9.2 | 22 | 80.7 ± 7.8 | 6 | | |

Table S3. Mean otolith increment widths for various hatch-month cohorts of juvenile Japanese sardine in the Kuroshio–Oyashio transitional region

| | Increment width (μm) by age in days | | | | | | | | | | | | | | | | | | | |
|---|---|---|---|---|---|---|---|---|---|---|---|---|---|---|---|---|---|---|---|---|
| | 10 d | | 20 d | | 30 d | | 40 d | | 50 d | | 60 d | | 70 d | | 80 d | | 90 d | | 100 d | |
| Hatch month | Mean ± SD | *n* | Mean ± SD | *n* | Mean ± SD | *n* | Mean ± SD | *n* | Mean ± SD | *n* | Mean ± SD | *n* | Mean ± SD | *n* | Mean ± SD | *n* | Mean ± SD | *n* | Mean ± SD | *n* |
| 2 | 3.2 ± 0.8 | 135 | 4.7 ± 1.0 | 135 | 6.4 ± 1.4 | 135 | 8.0 ± 1.7 | 135 | 9.7 ± 2.0 | 135 | 10.6 ± 1.7 | 135 | 10.4 ± 1.9 | 135 | 8.9 ± 2.1 | 131 | 7.2 ± 1.9 | 118 | 6.0 ± 1.7 | 67 |
| 3 | 4.4 ± 1.1 | 450 | 6.1 ± 1.4 | 450 | 7.7 ± 1.7 | 450 | 9.1 ± 2.2 | 449 | 10.2 ± 2.3 | 418 | 10.4 ± 2.3 | 294 | 9.0 ± 2.3 | 158 | 7.8 ± 2.1 | 83 | 6.8 ± 1.8 | 35 | 6.2 ± 1.4 | 20 |
| 4 | 4.5 ± 1.0 | 511 | 6.9 ± 1.4 | 511 | 8.3 ± 1.4 | 508 | 9.3 ± 1.6 | 402 | 9.4 ± 2.1 | 183 | 8.8 ± 2.7 | 79 | 8.4 ± 3.1 | 21 | | | | | | |
| 5 | 5.0 ± 1.0 | 81 | 7.9 ± 1.3 | 81 | 10 ± 1.9 | 80 | 9.9 ± 1.7 | 31 | | | | | | | | | | | | |

Table S4. Mean back-calculated standard lengths for various hatch-month cohorts of juvenile Japanese sardine in the Kuroshio–Oyashio transitional region

| | Back-calculated standard length (mm) by age in days | | | | | | | | | | | | | |
|---|---|---|---|---|---|---|---|---|---|---|---|---|---|---|
| | 40 d | | 50 d | | 60 d | | 70 d | | 80 d | | 90 d | | 100 d | |
| Hatch month | Mean ± SD | *n* | Mean ± SD | *n* | Mean ± SD | *n* | Mean ± SD | *n* | Mean ± SD | *n* | Mean ± SD | *n* | Mean ± SD | *n* |
| 2 | 27.3 ± 4.0 | 135 | 37.3 ± 5.5 | 135 | 49.0 ± 6.6 | 135 | 61.0 ± 7.1 | 135 | 71.7 ± 7.4 | 132 | 80.8 ± 8.1 | 121 | 86.5 ± 9.2 | 73 |
| 3 | 31.7 ± 4.7 | 465 | 42.4 ± 6.9 | 436 | 54.8 ± 9.3 | 311 | 67.6 ± 8.1 | 168 | 76.3 ± 7.7 | 89 | 84.6 ± 7.7 | 35 | 91.5 ± 5.1 | 21 |
| 4 | 32.9 ± 4.0 | 420 | 44.5 ± 5.8 | 203 | 56.6 ± 6.9 | 87 | 66.3 ± 7.6 | 24 | | | | | | |
| 5 | 35.6 ± 2.8 | 38 | | | | | | | | | | | | |
